# Entrainment effects and information processing in coupled oscillator models of auditory biomechanics

**DOI:** 10.1101/2025.09.09.675066

**Authors:** Bastian Epp, Mikkel Berrig Rasmussen

**Author notes:** (Electronic mail) (Bastian Epp).

## Abstract

The ability to process sound as a means of gaining information about the environment is shared by many species of animals. The auditory system of many vertebrates contains non-linear and active elements, hair cells, in the inner ear. These are considered elementary for high sensitivity, high selectivity, and wide dynamic range. However, the implications of the complex interplay between active hair cell function and inner ear mechanics on information processing are not yet fully understood. Here we use the basic properties of vertebrate ears to develop a numerical approach that captures active and nonlinear elements with the goal of quantifying effects of entrainment on information processing. We show that entrainment and the generation of clusters affect information processing, and that inclusion of these phenomena extends the scope of commonly used models of hearing. The results show ubiquitous phenomena that are robust against variation of model parameters. We show that the encoding of information emerges as a global effect in the system rather than locally in individual elements. These findings have implications for our understanding of tonotopy in, e.g., the peripheral auditory system, and link active and nonlinear dynamic behavior with information encoding in the inner ear. This will help the identification of the key principles that underlie hearing in complex acoustical environments with strict physiological restrictions.

The purpose of this study is to bridge entrainment effects with information processing in biological sensory systems. The results show the impact of self-organizing (spontaneous) and entrainment (externally driven) phenomena on the sensitivity, frequency selectivity, and information encoding capacity of the system. The frequency-specific sensitivity to external stimulation provides an explanation for physiological data recorded from the vertebrate auditory system. The link to information theory allows us to quantify the benefit of entrainment of sensory detection performance. Based on these results, acoustical, mechanical, and behavioral data can be evaluated within the same framework. This enables exploitation of the results for diagnostic and technological applications in, for example, audio signal processing.

## I. INTRODUCTION

The sense of hearing provides access to information encoded in sound. This information plays an elementary role in the detection of food or prey and in communication. With unprecedented performance, the healthy human auditory system enables communication in complex acoustical environments with multiple, simultaneously active sound sources (Bregman and McAdams, 1994). However, this ability is crucially dependent on the state of the inner ear. Even mild hearing damage leads to deficits in sound processing, often accompanied by the inability to separate sound sources in noisy environments (Graydon et al., 2019, for an overview). This is a sign of a reduced amount of information available to the auditory system. Some key properties of a healthy auditory system are well characterized. They include the detection of low-intensity sounds, the processing of a wide dynamic range of intensity, and high frequency selectivity (Hudspeth, 2008; Ashmore et al., 2010). Mammals, for example, are capable of detecting sounds with intensities that cause deflections in the range of nanometers in the inner ear structure. They are also capable of processing sound intensity spanning six orders of magnitude by making use of compressive nonlinearity. In addition to the high sensitivity and wide dynamic range, most species also show an extraordinary sharp tuning of the neural fibers that transmit sound information to the brain (Heil and Peterson, 2015). Some physiological components linked to these properties have been identified. In particular, high sensitivity and sharp tuning are attributed to the motility of specialized sensory cells (hair cells, HC) (Hudspeth, 2008; Ashmore et al., 2010). Changes in endocochlear potential, ototoxic drugs, or mechanical damage affect hair cell motility and cause reductions in sensitivity. The re-duction in sensitivity is often related to reductions in selectivity and, most importantly, ultimately challenges in communicating in complex acoustical environments. These findings suggest that hair cell motility acts as an essential, nonlinear active process, contributing to efficient extraction of information from sound.

The precise role of nonlinear dynamic aspects in hearing performance is, however, not yet fully understood. Individual hair bundles can be observed experimentally. They can spontaneously oscillate and show typical characteristics of an active oscillator tuned close to a bifurcation point, possibly to maximize sensitivity and selectivity (Dierkes et al., 2008). The intrinsic periodicity of a hair bundle can also be modified by an external stimulus (Levy et al., 2016), consistent with the phenomenon of entrainment of a limit cycle oscillator. The activity in the inner ear is related to the presence of otoacoustic emissions (OAEs) in or outside the ear canal found in many species (e.g. Manley, 2022; Bergevin et al., 2015). The generation mechanism of OAEs in the mammalian cochlea might be more complex compared to the oscillation of individual hair cells (Shera, 2003b, 2015). But the spontaneous activity in the inner ear is consistent with the presence of sound emitted spontaneously (spontaneous otoacoustic emissions, SOAEs) from many species (e.g. Manley, 2022; Bergevin et al., 2015). And OAEs also show strong correlations between different types (Bergevin et al., 2012), indicating common physical mechanisms. Interactions between SOAEs and external stimuli have been experimentally shown in a listening experiment in human listeners (Long, 1998). Listeners with indicators of sensory-neural hearing loss (and hence challenges in communicating in complex environments) generally do not show SOAEs and often have highly reduced other OAEs. This supports the idea that active, nonlinear function plays a crucial role in hearing (Moulin et al., 1991) or, on a more functional level, that cochlear amplification is essential for listening performance (Shera, 2015).

With the close connection of listening performance in complex environments, the state of the inner ear and the presence of OAEs, it becomes clear that efficient extraction and processing of information in the auditory system is highly dependent on the interplay of a number of, at least in parts, nonlinear and active elements.

A commonly used approximation to model sound processing in the inner ear is the description as a bank of independent, overlapping band-pass filters (e.g. Hohmann, 2002; Jepsen et al., 2008; Lyon, 2011). These models account for a variety of properties of local peripheral auditory processing, such as compression, different types of masking, and frequency selectivity. Increasing insights into inner ear physiology since the late 1960s (Guinan et al., 2012) indicate that some of the assumptions and limitations made in the filter-based approach ignore potentially important aspects of information processing for the sake of computational simplicity.

The recent developments in the observation and the mathematical description of the nonlinear dynamic properties of the hearing organ motivate an alternative approach to model sound processing in the peripheral auditory system. A model that can identify the ubiquitous and plausible dynamical phenomena present across species can shed light on the role of nonlinear dynamic properties in the process of hearing. Hence, with an outset in the vertebrate inner ear, the goal of the present study is to link the nonlinear dynamic effect of entrainment with information processing. An assumption of the model is that activity in the mechanical part of the vertebrate hearing organ can be modeled as a system of coupled, nonlinear and active oscillators. Based on previous findings of a) entrainment in coupled oscillators and b) the existence of active, nonlinear phenomena in physiological and behavioral data, we hypothesize that entrainment affects information encoding in the vertebrate inner ear. To this end, we quantify the dynamic phenomena that arise in a system of tonotopically arranged, coupled, nonlinear oscillators on various length- and time scales. We predict that entrainment in the undriven system leads to clustering, and that application of an external driving force will interfere with the system-intrinsic entrainment and lead to frequency-specific differences in susceptibility to the external driving force. Ultimately, if entrainment (and hence clustering phenomena) are beneficial for information processing, then the overall information of the driving force transferred into the system will be higher in the presence than in the absence of entrainment-related phenomena.

## II. MATERIAL AND METHODS

The numerical systems were implemented in JULIA (Bezanson et al., 2017). Each system consisted of *N* = 80 oscillators described by an individual equation of motion. The parameters were adjusted such that each oscillator in isolation displays a limit cycle in phase space. The intrinsic periodicity of the oscillators was adjusted so that the system of oscillators showed a gradient between the first and the last oscillator in the system. Coupling was introduced to the nearest neighbors, and the sinusoidal driving force *F*_*d*_(*t*) with frequency *f*_*d*_ and amplitude *a* was applied to all oscillators simultaneously.

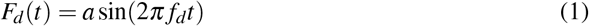

The first and last oscillators in the system were subject to free boundary conditions. The coupled set of equations was solved using the “tsit5” routine of JULIA (Tsitouras, 2011). The simulation results are shown after a simulated duration of 1500 s, excluding an oset phase of simulated 500 s.

The values of the model and coupling parameters were empirically chosen to obtain a similar number of clusters (see tab.I) for the different coupling conditions.

**TABLE I.** Parameter values of the coupled systems of van der Pol oscillators (VP) and Ginzburg Landau oscillators (GL).

|  | Van der Pol (VP) | Ginzburg Landau (GL) |
| --- | --- | --- |
| | $\{[\omega_1, \omega_{80}], \mu, d, k\}$ | $\{[\omega_1, \omega_{80}], \varepsilon, B, d, k\}$ |
| no coupling | $\{[1, 5] \cdot 2\pi, 0.5, 0, 0\}$ | $\{[1, 5] \cdot 2\pi, 4, 1, 0, 0\}$ |
| dissipative coupling | $\{[1, 5] \cdot 2\pi, 0.5, 15, 0\}$ | $\{[1, 5] \cdot 2\pi, 4, 1, 15, 0\}$ |
| reactive coupling | $\{[1, 5] \cdot 2\pi, 0.3, 0.15, 1\}$ | $\{[1, 5] \cdot 2\pi, 1, 1, 0.15, 1\}$ |

### A. Van der Pol model

One set of systems was based on van der Pol oscillators (VP), previously used to describe generic aspects of OAEs (but rejected as a specific model of mammalian cochlear mechanics) (e.g. Long and Talmadge, 1997). To ensure the same shape of the limit cycle for all oscillators in the system while modifying their periodicity, a modified version of the equation of motion was used (for details, see Sørensen et al., 2019):

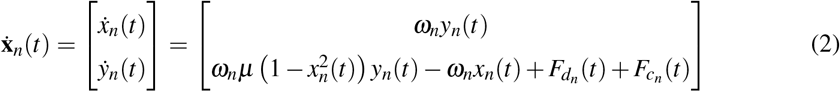

One set of equations was formulated for each oscillator *n* for amplitude *x*(*t*) and velocity 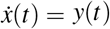. The parameter *µ* controlled the degree of nonlinearity. The value of *ω*_*n*_ controlled the intrinsic periodicity of each oscillator for an absent nonlinearity. 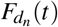 and 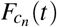 represent the external driving force and the force imposed by nearest-neighbor coupling, respectively.

Nearest-neighbor coupling was introduced by:

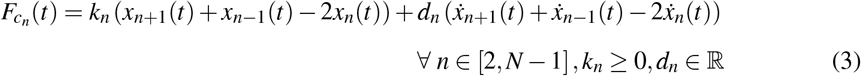

The parameters *k*_*n*_ and *d*_*n*_ allow to control the energy preserving (reactive) and energy dissipating (dissipative) coupling, respectively. The first (n=1) and last (n=N) oscillators in the system were only coupled to their next neighbor (2 and N-1, respectively).

### B. Vilfan-Duke model

The other set of systems was based on a simplified Ginzburg-Landau equation (GL), previously used to model Lizard SOAE (Vilfan and Duke, 2008). The equation of motion of the *n*-th oscillator was given by:

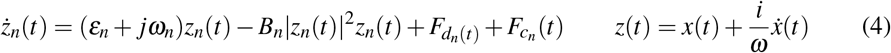

The real part of *z*_*n*_ can be interpreted as the amplitude *x*(*t*), and the imaginary part as proportional to the velocity 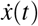 of the oscillator. The parameters *ε*_*n*_ and *B*_*n*_ control the speed of convergence to the limit cycle, the nonlinearity, and the radius of the phase portrait. The parameter *ω*_*n*_ controls the periodicity of the limit cycle. This type of oscillator shows a circular limit cycle in phase space. 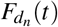 and 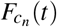 represent the external driving force and the force imposed by nearest-neighbor coupling, respectively. Nearest-neighbor coupling was implemented by:

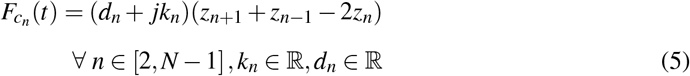

The parameters *k*_*n*_ and *d*_*n*_ allow to control the energy preserving (reactive) and energy dissipating (dissipative) coupling, respectively. The first (n=1) and last (n=N) oscillators in the system were only coupled to their next neighbor (2 and N-1, respectively).

### C. Estimation of instantaneous periodicity and clusters

To quantify the dynamics, the winding number *ρ* in phase space was used:

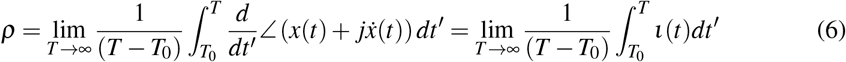

The kernel of the integral *ι*(*t*) will be used as a proxy measure of the instantaneous frequency of the oscillator. To avoid confusion with the basis functions of the Fourier domain, *ι*(*t*) will be referred to as “instantaneous periodicity” throughout this paper.

To gain information on the statistical moments of the distribution underlying the values of *ι*(*t*), values of *ι*(*t*) for all integration time steps and oscillators were stored. To compare the values of *ι*(*t*) with the driving force frequency, the results were normalized by 2*π*:

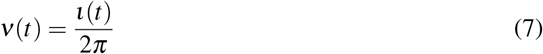

In the numerical calculation of average periodicity, the integral in eq.6 was replaced by the unwrapped phase at time *T* minus the unwrapped phase at time *T*_0_.

## D. Mutual information

Information in the oscillation of the system was quantified using mutual information with the unit of Shannon bits (Shannon, 1948):

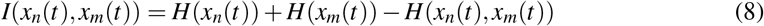

Where *H*(*x*_*n*_(*t*)) describes the entropy of signal *x*_*n*_(*t*) and *H*(*x*_*n*_(*t*), *x*_*m*_(*t*)) describes the joint entropy of the two signals *x*_*n*_(*t*) and *x*_*m*_(*t*):

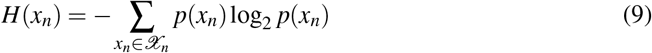

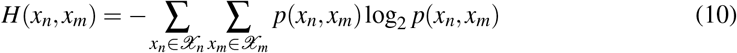

with *𝔛*_*n*_, *𝔛*_*m*_ representing the set of amplitudes in signals *x*_*n*_(*t*) and *x*_*m*_(*t*), respectively. *p*(*x*_*n*_, *x*_*m*_) represent the joint probability distribution of the amplitudes *x*_*n*_ and *x*_*m*_. Mutual information is always finite and quantifies the amount of information contained in both *x*_*n*_(*t*) and *x*_*m*_(*t*). In the tested systems, mutual information was quantified between all pairs of oscillators and between each oscillator and the external driving force.

To obtain an estimate of the overall mutual information between the driving force *F*_*d*_(*t*) and all *N* oscillators, the summed information was calculated by summation over all oscillators:

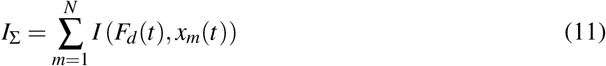

## III. RESULTS

### A. Clustering due to nearest-neighbor coupling

Figure 1 shows the state of two systems of coupled oscillators. Figure 1A-C shows systems of VP oscillators in the absence (A) and presence (B-C) of coupling, respectively. Figure 1D-F shows systems of GL oscillators in the absence (D) and presence (E-F) of coupling, respectively. In the absence of a driving force (A, D), the oscillators show an average periodicity that corresponds to the imposed intrinsic periodicity (dotted line). The VP oscillators (A) show a broad distribution of *ν* (red lines), in line with that VP oscillators show relaxation dynamic behavior and, as a consequence, do not have a circular phase portrait. The GL oscillators (D) show no variability in *ν*, in line with their harmonic oscillation behavior and their circular phase portrait.

**FIG. 1.**
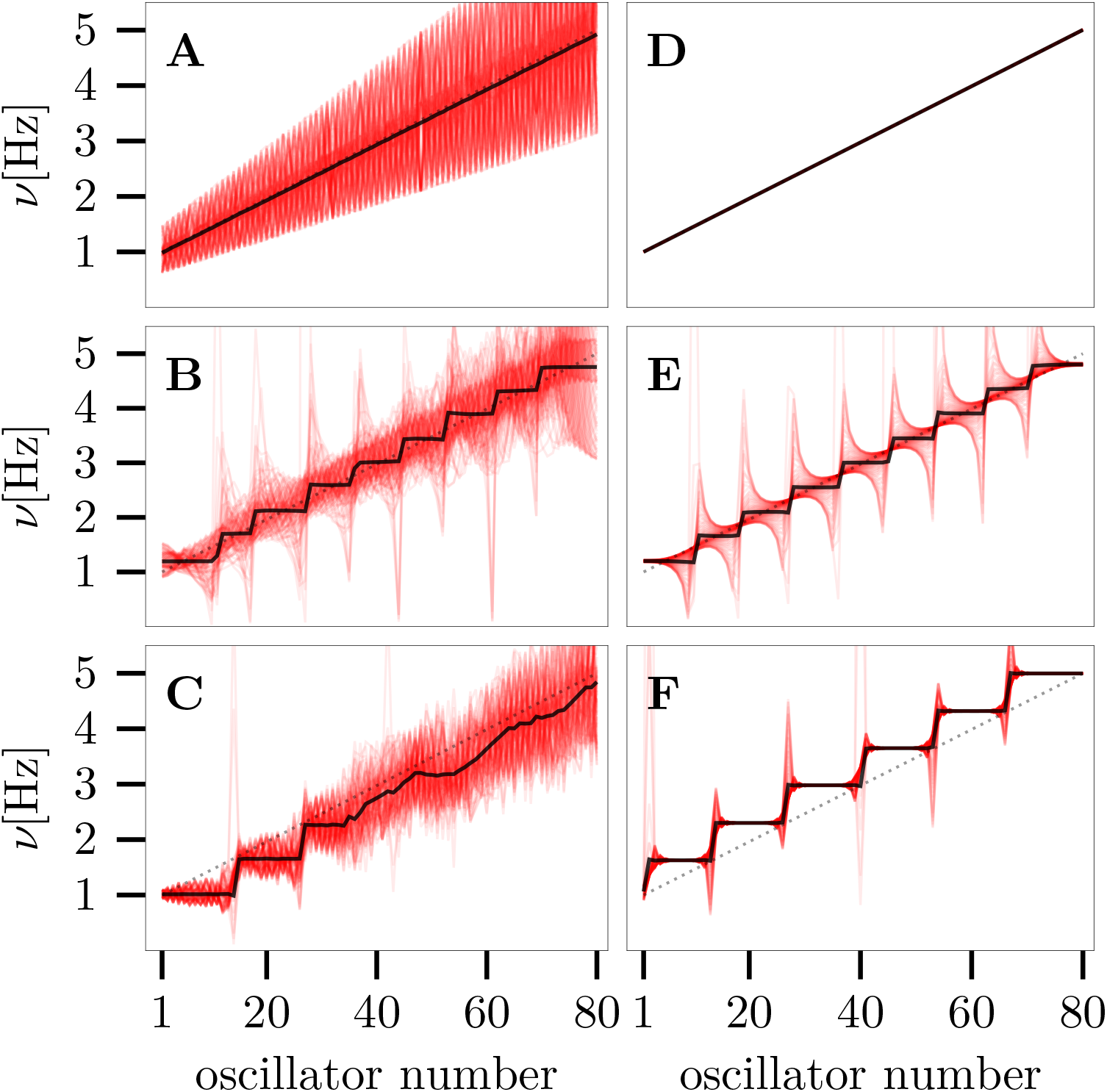
Systems of 80 coupled oscillators without external forcing. Panels A-C show systems composed of VP oscillators, panels D-F systems composed of GL oscillators. The first, second, and third rows shows systems without coupling (A,D), with dissipative coupling (B,E), and with dissipative- and reactive coupling (C,F). The dotted line shows the intrinsic periodicity of the oscillators. The solid line shows the average of the periodicity *ν* derived through the instantaneous periodicity *ι*. The red lines are a superposition of all values of 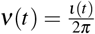 for each oscillator and time step.

With dissipative nearest-neighbor coupling (B, E), the oscillators in both systems organized into clusters of equal average periodicity 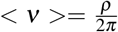 (solid black line), visible as plateaus (or “clusters”). For both systems, the clusters showed approximately equal size. The center of each cluster aligned approximately with the intrinsic periodicity of the corresponding oscillator. The variability of *ν* was higher for the VP (B) than for the GL (E) oscillators, consistent with the observation for uncoupled systems (A, D). In addition to the emergence of clusters, both systems showed a clearly systematic pattern of the distributions of *ν* within each cluster. Within each cluster, the distributions of *ν* became increasingly skewed towards the edges of the cluster, while the average periodicity stayed constant. Towards lower oscillator numbers (with lower intrinsic periodicity), the distributions became skewed towards higher periodicity. Towards higher oscillator numbers (with higher intrinsic periodicity), the distributions became skewed towards lower periodicity. Hence, the edges of each cluster bent away from the average periodicity of the neighbored cluster.

With reactive nearest-neighbor coupling (C, F), the oscillators organized, similarly to dissipative coupling, into clusters of equal periodicity. The VP (C) and the GL (F) system showed similar sizes of clusters, with the VP system showing fewer clusters. The VP system showed higher variability in *ν* than the GL system, consistent with the dissipative- and the uncoupled systems (A-B, D-E). The difference was the location of the clusters relative to the intrinsic periodicity of the oscillators. The clusters in the VP system (C) aligned with the intrinsic periodicity of the oscillator with the lowest number within each cluster, while the clusters in the GL system (F) aligned with the intrinsic periodicity of the oscillator with the highest number. This difference is the result of the sign of *k* in the system (see also Vilfan and Duke, 2008). For both systems, the distributions at the edges of the clusters were sharper with reactive coupling than with dissipative coupling and with smaller variation within each cluster.

On a long time scale, these results show a robust clustering of oscillators when coupled to their nearest neighbors, consistent with a previous model of lizard hearing (e.g. Vilfan and Duke, 2008). On shorter time scales and in addition to previous findings, the results also show a systematic variation of the instantaneous periodicity *ν* of each oscillator which depends on the specific position within a cluster, leading to systematic “tails” of *ν* at the cluster edges.

## B. Effect of driving force on clusters

Figure 2 shows the effect of a sinusoidal driving force (driving frequency of 3.5 Hz) on the clustering for the two systems. Figure 2A,B,C shows driven systems of VP oscillators with dissipative coupling. Figure 2D-I shows driven systems of GL oscillators with dissipative (D-F) and reactive (G-I) coupling. The driving amplitudes were low (*a* = 1; panels A, D, G), medium (*a* = 3; panels, B,E,H), and high (*a* = 5; panels C, F, I). The sinusoidal driving force resulted, consistently for all systems, in a modification of the cluster structure. With increasing driving force (Fig.2, rows from top to bottom), a cluster formed at the driving force frequency which included a higher number of oscillators than in the absence of the driving force. The sensitivity of the cluster arrangement on the external driving force depended on the oscillator type and the coupling parameters. The driving force also had an effect on the other oscillators and clusters in the system, even if their average periodicity was remote from that of the driving force. The addition of the sinusoidal driving force increased the variability of *ν* for the individual oscillators (compare to Fig.1). Also, the driving force lead to an asymmetry in the distributions of clusters: While clusters above the driving force frequency (higher oscillator numbers) tended to be preserved for the selected driving force amplitudes, clusters with lower periodicity tended to disappear with increased driving force amplitude. For certain combinations of system- and driving force parameters, the distributions of *ν* for oscillators within the largest cluster concentrated around narrowly distributed values (panel H, I), indicating a systematic interaction of the oscillator dynamic and the external driving force while keeping the average periodicity constant. Interesting to note is the co-existence of tails in the distributions of *ν* and clusters. Also in the cases where clusters were less pronounced indicated the presence of tails the presence of oscillator clustering (e.g., panel F).

**FIG. 2.**
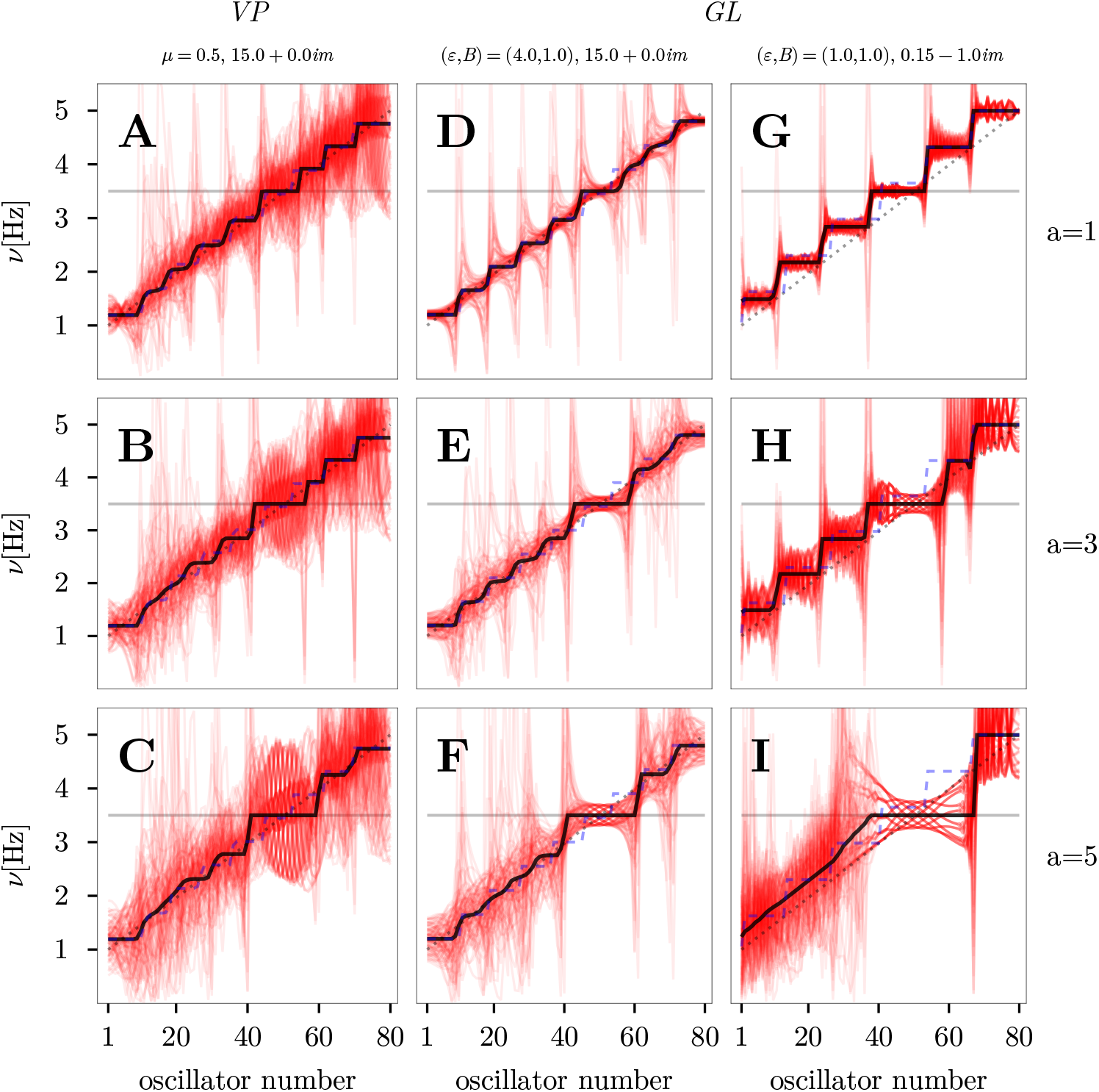
Same as fig. 1, but in the presence of external sinusoidal forcing (horizontal gray line) applied to all oscillators. Panels A-C show systems composed of VP oscillators with dissipative coupling. Panels D-F and G-H show systems composed of GL oscillators with dissipative and reactive coupling, respectively. The first, second, and third rows show the clustering with driving force amplitudes of 1, 3, and 5, respectively. The dashed blue line indicates the cluster structure of the corresponding undriven system (*a* = 0).

### C. Encoding of information in the absence and presence of a driving force

Figure 3 shows the mutual information of neighbored oscillators (heat map), and each individual oscillator with *F*_*d*_(*t*) (solid line). Results are shown for an uncoupled GL system (A-D), and the coupled GL systems from Fig. 2 (E-H, dissipatively coupled; I-L, reactively coupled).

**FIG. 3.**
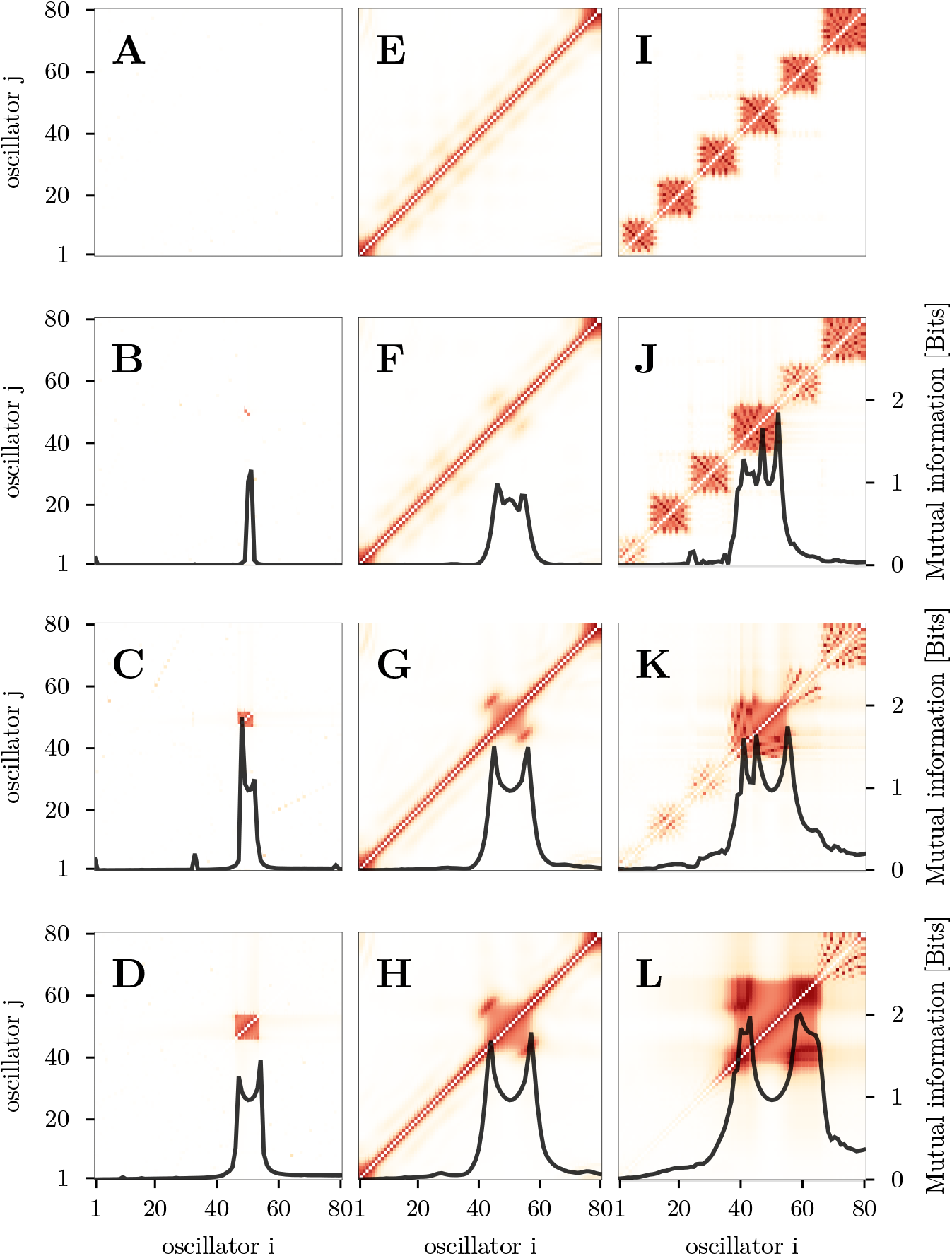
Mutual information of two neighbored oscillators (heat map, red) and with the external driving force (solid line, black) in systems of uncoupled (A-D) and coupled (E-L) GL oscillators. The coupled systems are the same as in fig. 2 D-F and G-I. The first row shows the undriven system. The second, third, and forth rows show the clustering with driving force amplitudes of 1, 3, and 5, respectively.

In the uncoupled system (A-D), mutual information only showed a limited structure across oscillators. For the lowest amplitude (B), the mutual information was generally low. With increased amplitude (C-D), the group of oscillators with high mutual information formed around the oscillator which had an intrinsic periodicity close to that corresponding to the driving force. This pattern reflected the susceptibility of each individual oscillator to adopt the external driving force periodicity. On a system level, this means that only a small group of elements encoded the information (frequency) in the driving signal. The mutual information with the driving force is restricted to a small number of oscillators, and shows a clear bi-modal distribution at the highest driving force amplitude (D).

For oscillators with dissipative (E-H) or reactive (I-L) coupling, the clustering of oscillators imposed a distinctively different pattern on the mutual information of neighbored oscillators compared to the uncoupled system. Due to the coupling, neighbored oscillators interacted and hence showed a high degree of mutual information. The addition of a driving force to a system with dissipative coupling affected this pattern by the emergence of a strongly connected area of oscillators within a cluster. The corners of the emerging structure are interesting to note, indicating higher mutual information at the edges of a cluster compared to the center of the cluster. It is also interesting to note that the other clusters in the system showed a similar, though weaker, pattern. For the investigated amplitudes and with dissipative coupling mainly the group of oscillators within the cluster aligned with the driving force frequency was affected by the driving force amplitude. The other oscillators were hardly affected. This was consistent with Fig.2D-F where the cluster structure was roughly conserved in the system with dissipative coupling. The mutual information of each oscillator with the external driving force was distributed over a higher number of oscillators compared to the uncoupled system and clearly showed a bimodal distribution.

In the system with reactive coupling (I-L), the addition of the driving force also changed the structure of mutual information in the system. For the lowest driving force amplitude (J), a dis-tinct structure was visible for groups of oscillators contained in one cluster. The higher mutual information in this system was consistent with a smaller amount of variability in *ν* in the oscillator clusters (see also Fig.2G). With increasing amplitude, the amount of mutual information was highest within the group of oscillators aligned with the driving force frequency. For the two highest amplitudes investigated (K, L), the driving force largely reduced the presence of mutual information in the other clusters. This is consistent with the increased variability within clusters (for the medium driving force amplitude, Fig.2H), and the absence of clusters for the highest driving force amplitude (Fig.2I). On system level, these results indicated a higher degree of encoding of the external driving force in the system compared to the uncoupled system. It is also interesting to note that the highest amount of mutual information was around the edges of the oscillator clusters. The mutual information of each oscillator with the driving force showed the broadest distribution of the investigated systems. Even at the lowest amplitude of the driving force, the information was present in a large number of oscillators, converging into a bimodal distribution at the highest amplitude of the driving force.

Figure 4 shows the summed mutual information *I*_Σ_ across all oscillators of a GL system in the presence of a driving force for a set of driving frequencies *f*_*d*_ from 0 to 7 Hz and driving amplitudes *a* from 0 to 10. Panels A-C show *I*_Σ_ for uncoupled, dissipatively coupled, and reactively coupled oscillators, respectively. *I*_Σ_ was low for low values of *a* and increased for increasing values of *a*. In the uncoupled system (A), *I*_Σ_ increased monotonically with *a*, relatively independent of *f*_*d*_. For driving frequencies below 1 Hz and above 5 Hz, high amplitudes were required to increase *I*_Σ_. For the coupled systems (B, C), lower amplitudes were required to reach the same value of *I*_Σ_ compared to the uncoupled system (A). Interesting to note are the vertical stripes visible in panels B and C. These local maxima of *I*_Σ_ reflect a high susceptibility of the system to the driving force when the frequency of the driving force coincided with a plateau of periodicity. A difference between panels B and C is the presence of a non-monotonic increase of *I*_Σ_ for low values of *a* (see also Figure 5). The data indicate a higher susceptibility to the driving force with amplitudes up to 1 compared to amplitudes between 1.2 and 2. The local increase in *I*_Σ_ was similar to the transition between the structures visible in Fig. 3 F-H and 3 J-L. While the structures differed between dissipative and reactive coupling for low amplitudes of the driving force (Fig.3F, J), the structures and hence the overall information present in the system converged for higher amplitudes of the driving force (Fig.3F, J).

**FIG. 4.**
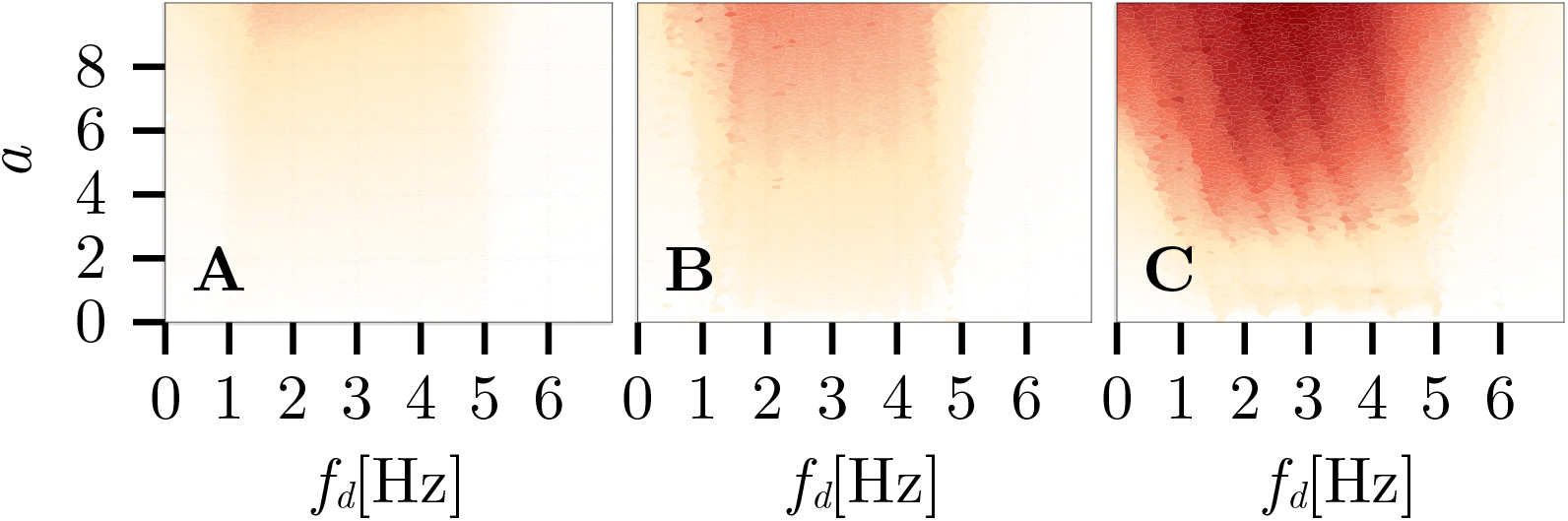
Summed mutual information *I*_Σ_ for the uncoupled (A), dissipatively coupled (B), and reactively coupled (C) GL oscillators shown in Figure 3. *I*_Σ_ was calculated for each combination of driving force amplitude *a* and driving force frequency *f*_*d*_. The plot shows normalized data.

**FIG. 5.**
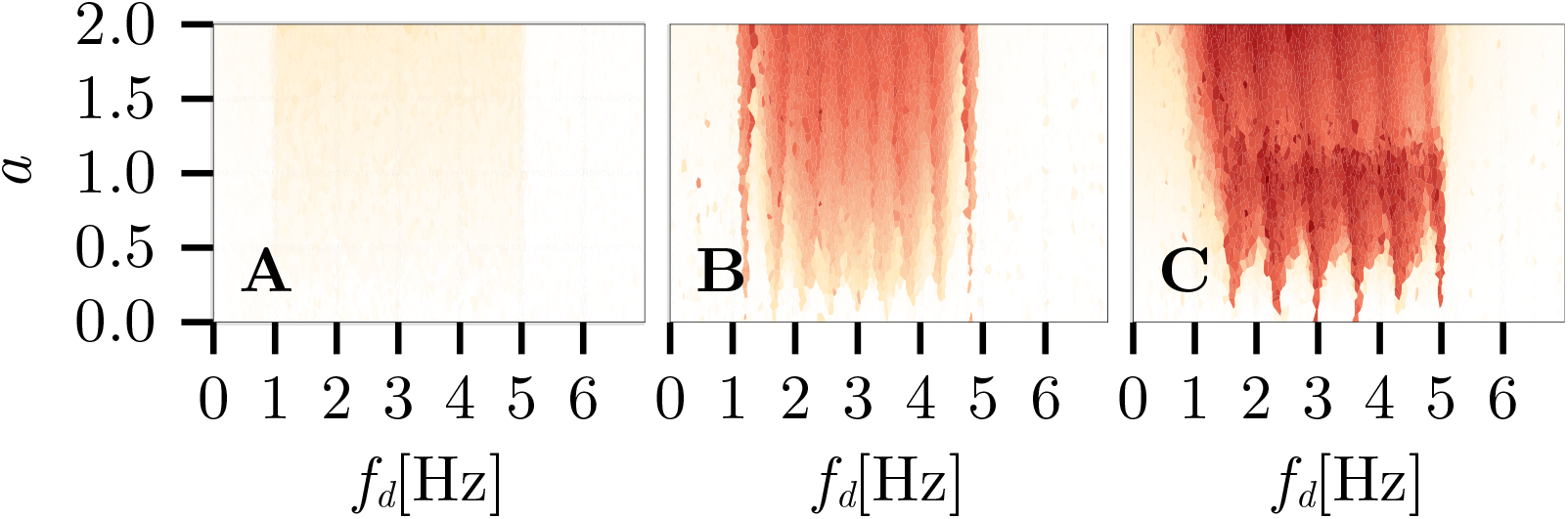
Same as Figure 4, but only for a subset of driving force amplitudes and higher resolution.

## IV. DISCUSSION

The simulations of the present study aimed to link mechanical processing with information processing of sound in the vertebrate inner ear. It is important to note that a direct comparison of the simulated systems is challenging because of their highly nonlinear nature and dynamic behavior. However, the presence of common phenomena provides relevant information about generalizable principles. The models in the present study were purely deterministic. The metrics used in the present study are usually not directly available from experimental data and should therefore be considered related but not identical to the metrics used in experimental studies. In addition, the impact of stochastic effects on the results is outside the scope of the present study but will need to be considered in the light of experimental data. In particular, the selected metric of “mutual information” only provides an upper bound of information, not considering redundancy or other implications imposed by the coupling of the individual oscillators. Hence, the observations need to be considered in a more generalized sense than in a system- and species-specific sense. The discussion will therefore focus on the common phenomena across the VP and the GL systems of oscillators.

## A. Comparison of simulated phenomena with experimental findings

The existence of spontaneous otoacoustic emissions (SOAE) has, among others, been linked to the presence of a dynamic behavior consistent with that of limit cycle oscillators (e.g. Talmadge et al., 1991; Vilfan and Duke, 2008; Wit et al., 2020). Properties that are consistent between SOAE and a limit cycle oscillator are, among others, the distribution of SOAE energy and spectral spacing in the frequency spectrum (e.g. Bialek and Wit, 1984; Talmadge et al., 1993; van Dijk and Wit, 1998). In the present study, a proxy to SOAE can be simulated by the superposition of the amplitudes of all oscillators. Figure 6 shows the magnitude spectrum of the simulated SOAE. The spectrum shows a series of narrow distributions of energy, very much like an SOAE spectrum in lizards or mammals.

**FIG. 6.**
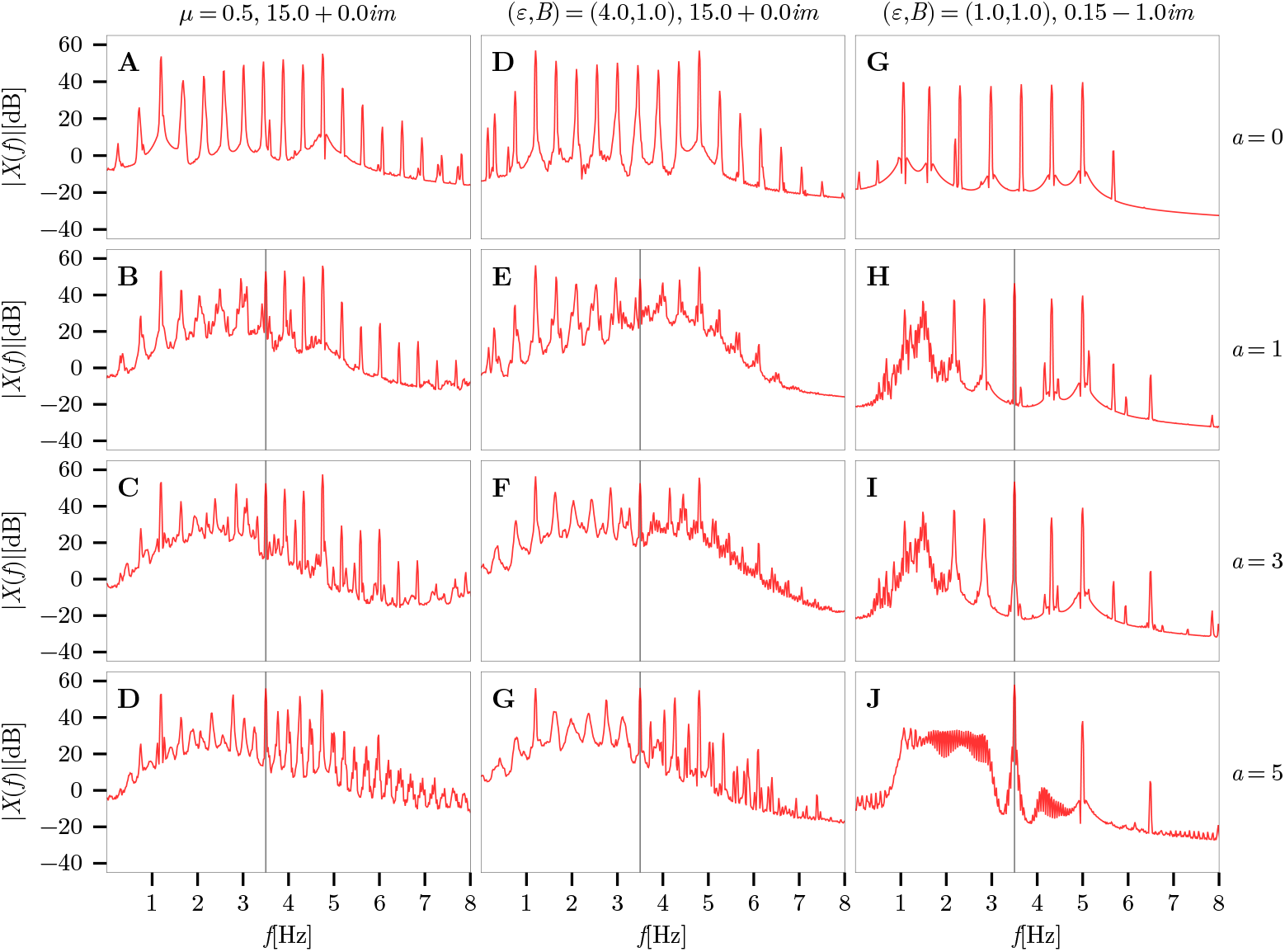
Magnitude spectrum of the summed time series of all oscillators for the systems and conditions shown in Figure 2. The vertical dashed lines indicate the driving force frequency.

Each peak in the frequency range of 1 to 5 Hz in this spectrum aligned with the frequency *ν* (derived as the inverse periodicity) of each cluster. The additional peaks are a result of the non-harmonic oscillatory behavior of the system. This similarity between simulated SOAE and frequency clusters supports the idea that SOAE can be the result of an interaction of active, non-linear oscillations in the inner ear leading to entrainment.

In particular, in agreement with the presence of an entrainment mechanism in the inner ear, the sudden transition between the presence and absence of beating between an SOAE and an external stimulus (Long, 1998) has been shown. In the framework of the present model, this transition corresponds to a change in the structure of the oscillator clusters as observed in Figure 2. Too weak stimuli will not be able to modify the periodicity of individual clusters, while stimulation with an increased amplitude is able to shift and even remove individual clusters. Under the assumption that a similar phenomenon is present in the inner ear, this is expressed as a change in the spectral distribution of the SOAE energy and the potential disappearance of individual peaks (see Figure 6). This dynamic is indeed in line with both theoretical and experimental results (van Dijk and Wit, 1990; Murphy et al., 1995a,b) and is also discussed in detail by Wit and Bell (2024).

Another interesting comparison can be made between the threshold microstructure and the alignment with SOAE (Zwicker and Schloth, 1984). The experimental data show a high sensitivity of the listener when the external stimulus coincides with the frequency of the SOAE in the magnitude spectrum of the ear canal signal. In terms of the present study, this can be interpreted in terms of the effect of an external stimulus on the clusters. In particular, in the case where the stimulation frequency coincides with the periodicity of a cluster, the external driving force will strengthen the impact of the cluster on the neighboring oscillators, increase its size, and reduce the number of periodicities in the system. One might speculate that a mechano-electrical transduction mechanism benefits from the oscillation resulting in the combination of self-sustained activity and external stimulation. In the opposite case where the driving frequency corresponds to a periodicity in between clusters, the oscillation imposed by the external stimulus needs to overcome the intrinsic clustering of the system and hence lead to a less efficient input into a mechano-electrical transduction mechanism. This interpretation is consistent with the presence of local maxima in the summed information *I*_Σ_ present in the coupled, but not the uncoupled system (Fig. 4).

For lizards, the findings can be related more directly. The GL oscillator system used in the present study has been proposed as a model for SOAE in lizards (Vilfan and Duke, 2008). Experimentally, individual hair bundles in the bullfrog have also been shown to oscillate spontaneously and can be synchronized by an external driving force (Levy et al., 2016). Models of individual hair bundles were successful in predicting key properties of dynamic behavior (e.g., Ó Maoiléidigh and Hudspeth, 2013; Dierkes et al., 2008; Roongthumskul and Bozovic, 2015). Hence, it is plausible that the creation of SOAE in lizards is related to an interaction of individually tuned oscillators that, in isolation, show intrinsic dynamic behavior but interact when embedded in a structure that allows for coupling. This argument is also in line with the finding that SOAEs are correlated across ears in species where the two ears are acoustically coupled (Roongthumskul et al., 2019).

An alternative explanation to the presence of individually coupled limit cycle oscillators is the concept of coherent reflections (Shera, 2003b). This method was used successfully to simulate a variety of phenomena, including SOAE (Epp et al., 2010). The present model approach is consistent with this idea when interpreting each limit cycle oscillator in the present model as a number of cochlear partitions that, driven by multiple reflections, behave like a limit cycle oscillator.

### B. Filtering by nonlinear dynamic processing

A common interpretation of the inner ear function is the filtering of the acoustic signal. Based on this, the most common models of auditory signal processing use some type of filter concept to describe this filtering process, partially explicitly assumed to be “cochlear filtering” (e.g. Hohmann, 2002; Jepsen et al., 2008; Lyon, 2011). Assuming a linear filtering, the stimulus would be filtered into separate “channels” that contain information within overlapping bands of frequency. In addition to the redundancy in the overlap region of the filters, these channels would be processed completely independently. Modifications of these filterbanks into level-dependent filters have some success in predicting data on interactions across frequency like two-tone suppression. However, common to all these models is the assumption of a static “characteristic frequency” to which a given channel is most sensitive. Here lies a fundamental difference between the use of digital filters and the model of coupled oscillators as implemented in the present study. Due to the nearest-neigbour coupling, the whole system of oscillators is mutually dependent. The nonlinear and active properties of each individual oscillator then leads to the observed collective phenomena like, e.g., clustering. This means effectively a shift in the “characteristic frequency” of a local “cochlear filter”, which is not possible with the most commonly used models based on digital filters. Also, the absence of coupling had large impact on the encodig of information between oscillators.

Both model approaches will be able to account for some elements of two-tone suppression, though based on different mechanisms. Nonlinear filter banks assume a shift in the operating point of the active process responsible for amplification. In coupled oscillator systems, the mechanism underlying the phenomenon of two-tone suppression can be linked to entrainment. A major difference between filterbank models and models including entrainment would be the information passed on the neural system. In filterbank models, overlapping filters will encode redundant information. Low-intensity frequency components might be present in the same channel as a nearby, higher-intensity frequency component. The neural system would then decide, based on some signal-to-noise criterion, which frequency would be relevant. In models including entrainment, the information passed to the neural system will not include such redundancy. If an oscillator is entrained to another periodicity, there will not be a “beating” due to simultaneous presence of two periodicities. Hence, the information passed on to neural “channels” will differ significantly from the filter-based models.

Although some phenomena can be explained by both approaches, coupled oscillators also include observed phenomena in OAE and perception (Long, 1998). On a more speculative basis, the effects of masking are consistent with entrainment and would make the assumption of an “internal signal-to-noise ratio” obsolete. An oscillator with high amplitude will affect the periodicity of neighboring oscillators and hence might entrain the distribution of periodicity within the whole system. Hence, the neural system would in this case only receive information about a single periodicity rather than two coexisting periodicities.

One of the main findings of the current study is the presence of “tails” in the distribution of *ν* within clusters of oscillators (Fig.1,2) and the connected maxima in mutual information (Fig.3). Assuming that the dynamic behavior of a specific oscillator is the input into a mechano-electric transduction mechanism, the response of the connected neuron will be different when connected to the middle or to an edge of a cluster. The average periodicity will be the same, and hence neural metrics quantifying average periodicity would likely not observe a difference. However, given the increased skewness of *ν* around the edges of the oscillator, the fine-grained neural information will differ. If that was the case, then the cluster with its “edges” will provide a high contrast between dominant periodicities in the system. This will also be true for driven systems. Assuming a connection between mutual information in the driven system and the information contained in the driving force, the simulation results show a higher degree of mutual information in the edges than in the middle of the clusters. This indicates an increased sensitivity of neurons that receive input from the edges of a cluster. Because entrainment, and hence clustering, is absent in the system of uncoupled oscillators, the higher amount of mutual information can be attributed to the presence of entrainment, facilitated by coupling.

## C. Implications for tonotopy

A key property of many mechanical inner ears is the presence of tonotopy. Experimentally, tonotopy is commonly measured using low-intensity pure-tone probes. The “characteristic place” (mechanical) or “characteristic frequency” (neural) is then defined as place of maximum amplitude for a given frequency or, equivalently, the frequency that causes maximum activity in a given neuron (e.g. Robles et al., 2008).

In the absence of entrainment and clustering, a mechanical system will show a clear and monotonic frequency-to-place mapping (Fig.1). In the presence of entrainment and clustering, the oscillations evoked by the external stimulus will interact with potentially present self-sustained oscillations. This interaction will lead to a re-organization of the coupled oscillators (Fig.2). For a weak external stimulus, the resulting “tonotopic” map will, to some extent, reflect the cluster structure of the system. The clusters will be visible as “plateaus” in the cases where the intrinsic oscillations of the system does not allow entrainment by the external stimulus. For high-intensity stimuli, the system response will be dominated by the intrinsic periodicity of each oscillator and show a monotonic transition of maximum response with frequency. This special case is consistent with experimental findings and the concept underlying filter banks.

A seemingly conflicting explanation for the staircase-like tonotopy has been given previously by Shera (2015) based on a different numerical model. In that model, the presence of multiple reflections caused by small variations in the strength of the model cochlear amplifier lead to a periodic variation of sensitivity along the modeled tonotopy. As a consequence, a staircase-like pattern of frequency sensitivity emerged. The model of Shera (2015) is built on a quasi-linear approach, implementing level-specific impedance values of stable coupled oscillators. This is in contrast to the limit-cycle oscillators of the present study, as well as models of the lizard ear (Vilfan and Duke, 2008). But these two approaches can be mapped onto each other: The model of Shera (2015) finds a staircase-like pattern as an emergent effect caused by multiple coherent reflections with low-intensity stimulation. One can assume that a suitable choice of parameters can lead to self-sustained activity in this model, also in the absence of a driving force. A model of coupled limit cycle oscillators, on the other hand, leads to the emergent effect of frequency clusters. These can be interpreted as self-sustained activity in the absence of external stimulation (SOAE in experimental terms), and lead to a similar staircase-like pattern of sensitivity to external stimulation. Hence, the self-sustained activity postulated in Shera (2015) as an emergent effect can be interpreted as the self-organized state of a set of limit cycle oscillators. An important difference between these two explanations lies in the boundary conditions. In the model by Shera (2015), the amount energy reflection back into the cochlea is an important parameter to achieve the selective sensitivity. A model of coupled limit cycle oscillators does not depend on a mechanism of multiple reflection.

Overall it might be expected that the mapping between “frequency” and “place” differs significantly in uncoupled (filter-based) and coupled systems when exposed to stimuli that are more complex than pure tones. Hence, the encoding of ecologically valid sounds will depend more on the information received by the neurons after mechano-electrical transduction than on the tonotopic organization of the system in the presence of pure tones of low intensity.

## V. CONCLUSION

Coupling of limit-cycle oscillators leads to entrainment-related clustering effects with “tails” in the distribution of the instantaneous frequency that affect temporal coding. Clustering in combination with a driving force leads to an amplitude- and coupling-dependent distribution of information across oscillators. The presence of clusters modulates the susceptibility of the system to the frequency of the external driving force. Hence, entrainment effects in coupled, spontaneously active nonlinear systems of oscillators lead to emergent, global information encoding and lead to frequency-specific differences in sensitivity to external stimulation. Applied as a model of vertebrate inner ears, this approach allows us to link inner ear mechanics, otoacoustic emissions and behavioral data to identify the biophysical mechanisms underlying high sensitivity, high selectivity, and wide dynamic range.

## VI. ACKNOWLEDGEMENTS

## REFERENCES

Ashmore, J., Avan, P., Brownell, W.E., Dallos, P., Dierkes, K., Fettiplace, R., Grosh, K., Hackney, C.M., Hudspeth, a.J., Jülicher, F., Lindner, B., Martin, P., Meaud, J., Petit, C., Santos Sacchi, J.R., Canlon, B., 2010. The remarkable cochlear amplifier. Hearing Research 266, 1–17. URL: http://dx.doi.org/10.1016/j.heares.2010.05.001, doi:doi: 10.1016/j.heares.2010.05.001.

Bergevin, C., Fulcher, A., Richmond, S., Velenovsky, D., Lee, J., 2012. Interrelationships between spontaneous and low-level stimulus-frequency otoacoustic emissions in humans. Hearing Research 285. doi:doi:10.1016/j.heares.2012.02.001.

Bergevin, C., Manley, G.A., Köppl, C., 2015. Salient features of otoacoustic emissions are common across tetrapod groups and suggest shared properties of generation mechanisms. Proceedings of the National Academy of Sciences 112, 3362–3367. URL: http://www.pnas.org/lookup/doi/10.1073/pnas.1418569112, xdoi:doi:10.1073/pnas.1418569112.

Bezanson, J., Edelman, A., Karpinski, S., Shah, V.B., 2017. Julia: A fresh approach to numerical computing. SIAM Review 59, 65–98. URL: https://epubs.siam.org/doi/10.1137/141000671, xdoi:doi:10.1137/141000671.

Bialek, W., Wit, H.P., 1984. Quantum limits to oscillator stability: Theory and experiments on acoustic emissions from the human ear. Physics Letters A 104, 173–178. doi:doi:10.1016/0375-9601(84)90371-2.

Bregman, A.S., McAdams, S., 1994. Auditory Scene Analysis: The Perceptual Organization of Sound. The Journal of the Acoustical Society of America 95. doi:doi:10.1121/1.408434.

Dierkes, K., Lindner, B., Jülicher, F., 2008. Enhancement of sensitivity gain and frequency tuning by coupling of active hair bundles. Proceedings of the National Academy of Sciences of the United States of America 105, 18669–18674. URL: https://www.pnas.org/doi/abs/10.1073/pnas.0805752105, xdoi:doi:10.1073/PNAS.0805752105/ASSET/0B0D45CD-ABA5-4F0E-AC37-07DB2525C231/ASSETS/GRAPHIC/ZPQ9990855100005.JPEG.

van Dijk, P., Wit, H.P., 1990. Synchronization of spontaneous otoacoustic emissions to a 2 f1-f2 distortion product. The Journal of the Acoustical Society of America 88. doi:doi: 10.1121/1.399734.

van Dijk, P., Wit, H.P., 1998. Correlated amplitude fluctuations of spontaneous otoacoustic emissions. The Journal of the Acoustical Society of America 104. doi:doi:10.1121/1.423259.

Epp, B., Verhey, J.L., Mauermann, M., 2010. Modeling cochlear dynamics: interrelation between cochlea mechanics and psychoacoustics. J Acoust Soc Am 128, 1870–83. URL: http://www.ncbi.nlm.nih.gov/pubmed/20968359, xdoi:doi:10.1121/1.3479755.

Graydon, K., Waterworth, C., Miller, H., Gunasekera, H., 2019. Global burden of hearing impairment and ear disease. Journal of Laryngology and Otology 133. doi:doi: 10.1017/S0022215118001275.

Guinan, J.J., Salt, A., Cheatham, M.A., 2012. Progress in cochlear physiology after Békésy. doi:doi:10.1016/j.heares.2012.05.005.

Heil, P., Peterson, A.J., 2015. Basic response properties of auditory nerve fibers: a review. doi:doi: 10.1007/s00441-015-2177-9.

Hohmann, V., 2002. Frequency analysis and synthesis using a Gammatone filterbank. Acta Acustica united with Acustica 88, 433–442.

Hudspeth, A.J., 2008. Making an Effort to Listen: Mechanical Amplification in the Ear. Neuron 59, 530–545. doi:doi:10.1016/j.neuron.2008.07.012.

Jepsen, M.L., Ewert, S.D., Dau, T., 2008. A computational model of human auditory signal processing and perception. The Journal of the Acoustical Society of America 124. doi:doi: 10.1121/1.2924135.

Levy, M., Molzon, A., Lee, J.H., Kim, J.W., Cheon, J., Bozovic, D., 2016. High-order synchronization of hair cell bundles. Scientific Reports 6. doi:doi:10.1038/srep39116.

Long, G., 1998. Perceptual consequences of the interactions between spontaneous otoacoustic emissions and external tones. I. Monaural diplacusis and aftertones. Hearing Research 119, 49–60. doi:doi:10.1016/S0378-5955(98)00032-X.

Long, G., Talmadge, C., 1997. Spontaneous otoacoustic emission frequency is modulated by heartbeat. The Journal of the Acoustical Society of … 102, 2831–2848. URL: http://link.aip.org/link/?JASMAN/102/2831/1.

Lyon, R.F., 2011. Cascades of two-pole-two-zero asymmetric resonators are good models of peripheral auditory function. J Acoust Soc Am 130, 3893–904. URL: http://www.ncbi.nlm.nih.gov/pubmed/22225045, xdoi:doi:10.1121/1.3658470.

Manley, G.A., 2022. Otoacoustic Emissions in Non-Mammals. Audiology Research 12, 260–272.

Moulin, A., Collet, L., Delli, D., Morgon, A., 1991. Spontaneous otoacoustic emissions and sensori-neural hearing loss. Acta Oto-Laryngologica 111. doi:doi: 10.3109/00016489109138419.

Murphy, W.J., Talmadge, C.L., Tubis, A., Long, G.R., 1995a. Relaxation dynamics of spontaneous otoacoustic emissions perturbed by external tones. I. Response to pulsed single-tone suppressors. The Journal of the Acoustical Society of America 97, 3702–3710. URL: http://asa.scitation.org/doi/10.1121/1.412387, xdoi:doi:10.1121/1.412387.

Murphy, W.J., Tubis, A., Talmadge, C.L., Long, G.R., 1995b. Relaxation dynamics of spontaneous otoacoustic emissions perturbed by external tones. II. Suppression of interacting emissions. The Journal of the Acoustical Society of America 97, 3711–3720. URL: http://asa.scitation.org/doi/10.1121/1.412388, xdoi:doi:10.1121/1.412388.

Ó Maoiléidigh, D., Hudspeth, A.J., 2013. Effects of cochlear loading on the motility of active outer hair cells. Proceedings of the National Academy of Sciences of the United States of America 110. doi:doi:10.1073/pnas.1302911110.

Robles, L., Robles, L., Ruggero, M.a., Ruggero, M.a., 2008. Luis Robles and Mario A. Ruggero. Physiological Reviews, 1305–1352.

Roongthumskul, Y., Bozovic, D., 2015. Mechanical amplification exhibited by quiescent saccular hair bundles. Biophysical Journal 108. doi:doi:10.1016/j.bpj.2014.11.009.

Roongthumskul, Y., Ó Maoiléidigh, D., Hudspeth, A.J., 2019. Bilateral Spontaneous Otoacoustic Emissions Show Coupling between Active Oscillators in the Two Ears. Biophysical Journal 116. doi:doi:10.1016/j.bpj.2019.02.032.

Shannon, C.E., 1948. A mathematical theory of communication. The Bell System Technical Journal 27, 379–423. doi:doi:10.1002/j.1538-7305.1948.tb01338.x.

Shera, C.A., 2003b. Mammalian spontaneous otoacoustic emissions are amplitude-stabilized cochlear standing waves. The Journal of the Acoustical Society of America 114, 244–262. doi:doi:10.1121/1.1575750.

Shera, C.A., 2015. The Spiral Staircase: Tonotopic Microstructure and cochlear Tuning The Journal of Neuroscience 35, 4683–4690.

Sørensen, L.M., Bysted, P.L., Epp, B., 2019. Clustering in an array of nonlinear and active oscillators as a model of spontaneous otoacoustic emissions. Proceedings of the International Congress on Acoustics 2019-Septe, 688–695. doi:doi:10.18154/RWTH-CONV-239452.

Talmadge, C.L., Long, G.R., Murphy, W.J., Tubis, a., 1993. New off-line method for detecting spontaneous otoacoustic emissions in human subjects. Hear Res 71, 170–82. URL: http://www.ncbi.nlm.nih.gov/pubmed/8113135.

Talmadge, C.L., Tubis, a., Wit, H.P., Long, G.R., 1991. Are spontaneous otoacoustic emissions generated by self-sustained cochlear oscillators? The Journal of the Acoustical Society of America 89, 2391–9. URL: http://www.ncbi.nlm.nih.gov/pubmed/1860998, doi:doi: 10.1121/1.400958.

Tsitouras, C., 2011. Runge-Kutta pairs of order 5 (4) satisfying only the first column simplifying assumption. Computers & Mathematics with Applications 62, 770–775.

Vilfan, A., Duke, T., 2008. Frequency clustering in spontaneous otoacoustic emissions from a lizard’s ear. Biophysical Journal 95, 4622–4630.

Wit, H.P., Bell, A., 2024. Something in Our Ears Is Oscillating, but What? A Modeller’s View of Efforts to Model Spontaneous Emissions. doi:doi:10.1007/s10162-024-00940-7.

Wit, H.P., Manley, G.A., van Dijk, P., 2020. Modeling the characteristics of spontaneous otoacoustic emissions in lizards. Hearing Research 385. doi:doi:10.1016/j.heares.2019.107840.

Zwicker, E., Schloth, E., 1984. Interrelation of different oto-acoustic emissions. The Journal of the Acoustical Society of America 75, 1148–1154. URL: http://link.aip.org/link/jasman/v75/i4/p1148/s1.

